# Functional interrogation of cellular Lp(a) uptake by genome-scale CRISPR screening

**DOI:** 10.1101/2024.05.11.593568

**Authors:** Taslima G. Khan, Juliana Bragazzi Cunha, Chinmay Raut, Michael Burroughs, Sascha N. Goonewardena, Alan V. Smrcka, Elizabeth K. Speliotes, Brian T. Emmer

## Abstract

An elevated level of lipoprotein(a), or Lp(a), in the bloodstream has been causally linked to the development of atherosclerotic cardiovascular disease and calcific aortic valve stenosis. Steady state levels of circulating lipoproteins are modulated by their rate of clearance, but the identity of the Lp(a) uptake receptor(s) has been controversial. In this study, we performed a genome-scale CRISPR screen to functionally interrogate all potential Lp(a) uptake regulators in HuH7 cells. Strikingly, the top positive and negative regulators of Lp(a) uptake in our screen were *LDLR* and *MYLIP*, encoding the LDL receptor and its ubiquitin ligase IDOL, respectively. We also found a significant correlation for other genes with established roles in LDLR regulation. No other gene products, including those previously proposed as Lp(a) receptors, exhibited a significant effect on Lp(a) uptake in our screen. We validated the functional influence of LDLR expression on HuH7 Lp(a) uptake, confirmed *in vitro* binding between the LDLR extracellular domain and purified Lp(a), and detected an association between loss-of-function *LDLR* variants and increased circulating Lp(a) levels in the UK Biobank cohort. Together, our findings support a central role for the LDL receptor in mediating Lp(a) uptake by hepatocytes.

## INTRODUCTION

Lipoprotein(a), or Lp(a), was discovered in 1963 as a unique variant of low-density lipoprotein (LDL)^1^. Early cross-sectional and retrospective studies suggested an association between elevated Lp(a) and coronary artery disease (CAD)^2,3^ that was confirmed in multiple larger prospective studies^4,5^. Lp(a) levels vary from ˂1 to ˃200 mg/dL in the general population, with each 2-fold increase associated with an estimated 22% greater risk of myocardial infarction^6,7^. Human genetic studies have established the causal influence of Lp(a) elevation on the development of both atherosclerotic cardiovascular disease^7–12^ and calcific aortic valve stenosis^13–16^, a common cause of heart failure and death. In light of this evidence, Lp(a) has become an attractive target for therapeutic development. While a clinical trial for *LPA* antisense oligonucleotide-based therapy is eagerly anticipated^17^, clinical experience with LDL-targeted treatments has demonstrated the benefits of a multifaceted approach to lowering atherogenic lipoproteins.

Lp(a) levels are highly heritable and largely determined by heterogeneity at the *LPA* locus, which encodes the apolipoprotein(a), or apo(a), component of Lp(a)^18,19^. Copy number variants in the KIV2 domain of *LPA* are inversely correlated with Lp(a) levels^20^, presumably due to more efficient maturation and secretion of smaller apo(a) isoforms^21,22^. The steady state level of circulating Lp(a) also depends on its rate of clearance, which occurs primarily in the liver^23,24^. The molecular basis of Lp(a) clearance, however, remains poorly understood. Multiple studies of the LDL receptor (LDLR) in cells^25–32^, animal models^23,30,33,34^, and humans^8,26,35–46^ have provided inconsistent or conflicting results. Over time, several other candidate Lp(a) receptors have been proposed, including other lipoprotein receptors^47,48^, toll-like receptors^49^, scavenger receptors^50,51^, lectins^24,52^, and plasminogen receptors^27,31^. It has also been proposed that multiple receptors may contribute to Lp(a) clearance^53^. In general, prior functional studies of Lp(a) uptake have involved a variety of cell types and methods for Lp(a) purification and detection. To date, no study has reported a systematic and unbiased functional interrogation of all potential Lp(a) receptors in the same experimental context.

We previously applied a genome-scale CRISPR screen to successfully identify regulators of cellular LDL endocytosis^54^. We readily detected expected genes encoding the LDL receptor and its canonical transcriptional and posttranscriptional regulators. We also detected novel regulators of LDL uptake that were reproducible across independent replicates and during follow up testing^54,55^. Given the excellent technical performance of this approach, in this study we sought to adapt it to identify functional modifiers of Lp(a) uptake. The results of our screen and our subsequent validation, mechanistic investigation, and analysis of human genetic variants all support a primary role for the LDL receptor in mediating Lp(a) uptake by hepatocytes.

## RESULTS

### Development of a flow cytometry-based assay for cellular Lp(a) uptake

To enable a forward genetic approach, we first set out to develop a method for the sensitive and specific selection of individual cells with aberrant Lp(a) uptake. Our initial attempts to fluorescently label purified Lp(a) preparations with either NHS-ester conjugation of primary amines or lipid labeling with DiI were complicated by sample precipitation, which was apparent both by gross visual inspection (Fig 1A) and by SDS-PAGE showing an inability of apolipoproteins to enter the stacking gel (Fig 1B). This effect was not observed in parallel labeling reactions of LDL, suggesting that the apo(a) component of Lp(a) sensitized samples to precipitation. In an attempt to reduce the hydrophobicity of our labeling approach, we next tested the fluorophore pHrodo iFL Red, which utilizes an STP-ester amine-labeling strategy with increased water solubility^56^. This approach circumvented lipoprotein precipitation, as labeled samples exhibited the same solubility and electrophoretic mobility as unlabeled samples (Fig 1C). After pHrodo iFL Red labeling and extensive dialysis to remove free dye, we recovered >90% of the original Lp(a) preparation with no disruption of sample purity or disulfide linkage between apo(a) and apolipoprotein B, as reflected by a shift in protein electrophoretic mobility under nonreducing conditions (Fig 1C). Incorporation of fluorophore into Lp(a) was confirmed by in-gel fluorescent scanning (Fig 1C) and by flow cytometry revealing a highly sensitive, time-dependent, and dose-dependent uptake of labeled Lp(a) by HuH7 cells (Fig 1D-E). Consistent with receptor-mediated endocytosis, co-incubation with a molar excess of unlabeled Lp(a) led to a dose-dependent competitive inhibition of fluorescent Lp(a)uptake (Fig 1F). Together, these findings support the suitability of our fluorescent Lp(a) labeling and cellular uptake strategy for high-throughput genetic screening.

**Figure 1.**
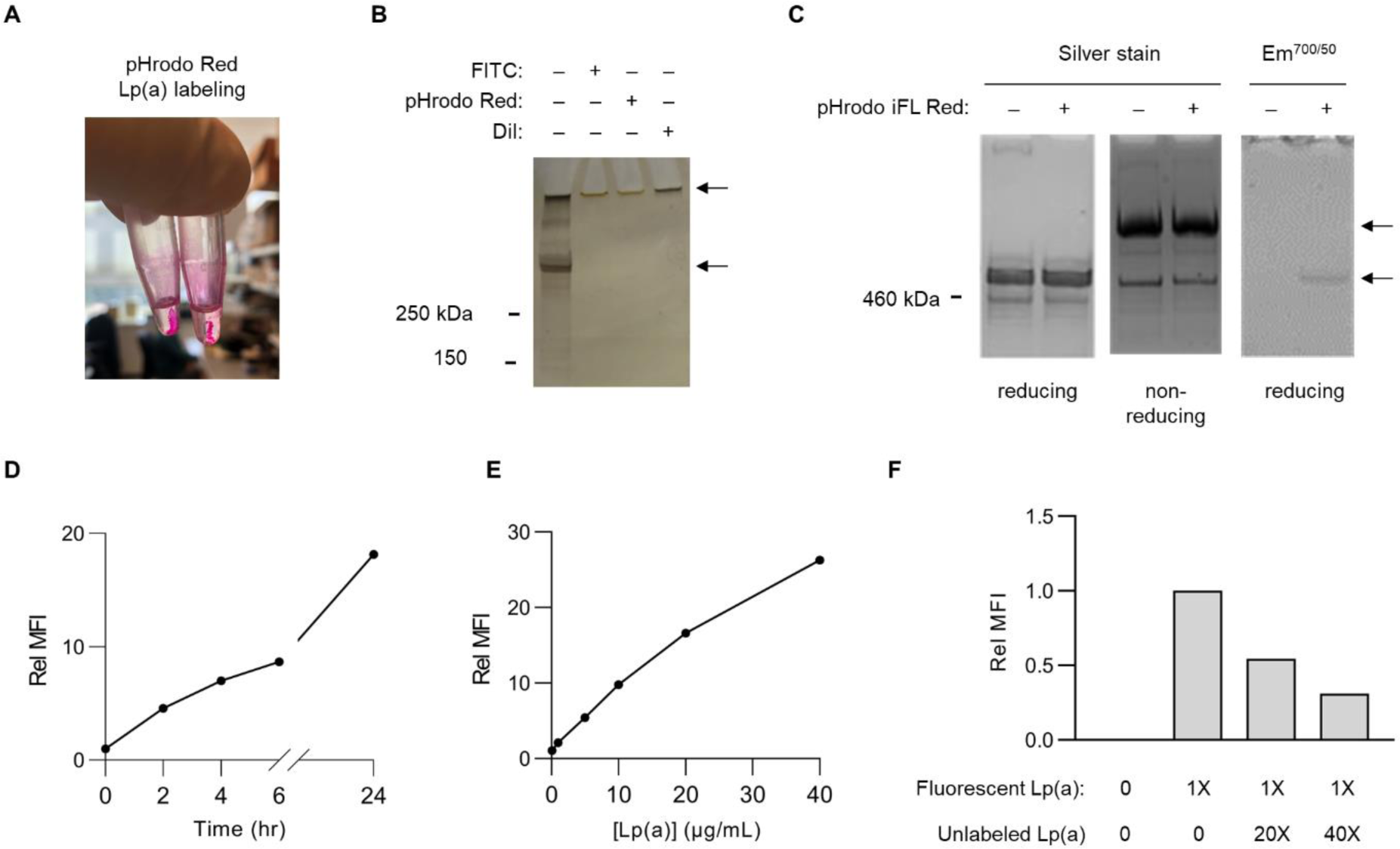
Development of a method for fluorescent labeling of purified Lp(a) and quantification of its cellular uptake with flow cytometry. (A) Overt sample precipitation during attempted fluorescent labeling of purified Lp(a) preparations with pHrodo Red. (B) SDS-PAGE and silver straining of purified Lp(a) preparations before and after attempted fluorescent labeling with FITC, pHrodo Red, and DiI. Upper arrow corresponds to precipitated protein remaining in loading well. Lower arrow corresponds to expected size (∼500 kDa) of co-migrating apo(a) and apolipoprotein B proteins. (C) SDS-PAGE under reducing and nonreducing conditions of a purified Lp(a) preparation before and after fluorescent labeling with pHrodo iFL Red followed by silver staining or in-gel fluorescence scanning. Upper arrow corresponds to expected migration of disulfide-linked apo(a)-apolipoprotein B complexes under nonreducing conditions; lower arrow corresponds to expected comigration of noncomplexed apo(a) and apolipoprotein B under reducing conditions. (D) Time course analysis of HuH7 cellular uptake of 10 µg/mL pHrodo iFL Red-labeled Lp(a). (E) Concentration-dependence of HuH7 cellular uptake of pHrodo iFL Red-labeled Lp(a) over 2 hrs. (F) Dose-dependent competitive inhibition of HuH7 cellular fluorescent Lp(a) uptake by co-incubation with a molar excess of unlabeled Lp(a). Representative flow cytometry plots are provided in Supplemental Figure 1.

### Genome-scale CRISPR screen for Lp(a) uptake regulators

Informed by the optimization experiments described above, we next scaled our cellular Lp(a) uptake assay to query the same GeCKOv2 genome-wide CRISPR knockout library^57^ we tested in our prior screen of LDL uptake^54^ (Fig 2A). For each of 3 independent biologic replicates, we transduced ∼60 million HuH7 cells at multiplicity of infection (MOI) ∼0.5, passaged cells for 14 days to allow for target site editing and turnover of residual protein, incubated pools of edited cells with fluorescent Lp(a), sorted 10% subpopulations of cells with the least and greatest Lp(a) uptake, and quantified gRNA abundance in each sample by massively parallel sequencing.

**Figure 2.**
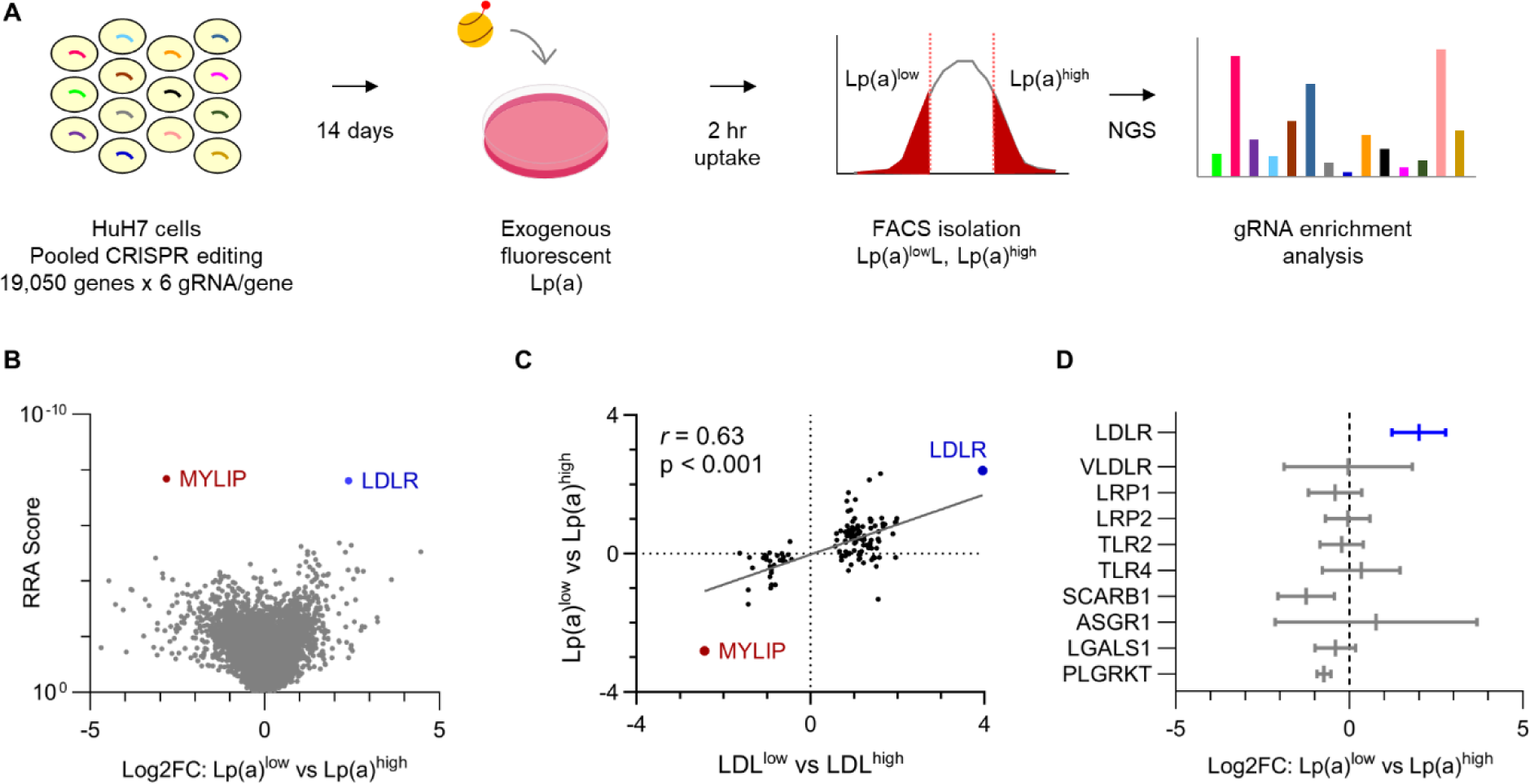
A genome-scale CRISPR screen for modifiers of HuH7 cellular Lp(a) uptake. (A) Schematic overview of screening strategy, with pools of HuH7 cells transduced with the GeCKOv2 genome-wide CRISPR knockout library at low MOI, passaged for 14 days, incubated with fluorescent Lp(a) for 2 hrs, and sorted into Lp(a)^low^ and Lp(a)^high^ subpopulations for extraction of genomic DNA and quantification of individual gRNA abundance by next-generation sequencing (NGS). (B) Volcano plot of aggregate gene-level Robust Rank Aggregation (RRA) scores (y-axis) relative to log2 fold change of normalized gRNA counts in Lp(a)^low^ relative to Lp(a)^high^ subpopulations. Genes whose disruption was associated with a significant (FDR<5%) decrease or increase in Lp(a) uptake are labeled and highlighted in blue and red, respectively. (C) Analysis of genes previously identified in our analogous prior screen of cellular LDL uptake, with the aggregate log2 fold change for gRNAs targeting each gene in LDL^low^ relative to LDL^high^ cells (x-axis) in that screen plotted relative to gRNA log2 fold change in Lp(a)^low^ relative to Lp(a)^high^ cells in this study. (D) Mean and 95% confidence intervals of aggregate gRNA log2 fold enrichment in Lp(a)^low^ relative to Lp(a)^high^ cells across 3 independent biologic replicates for genes encoding receptors previously proposed as mediators of cellular Lp(a) uptake. RRA scores, aggregate log2 fold change, and FDR values were calculated by MAGeCK; 95% confidence intervals of independent replicates were calculated using GraphPad Prism. Source data are provided in Supplemental Tables 1 and 2.

Analysis of gRNA enrichment across screen replicates revealed that the most significant decrease in Lp(a) uptake was conferred by disruption of *LDLR* while the most significant increase was conferred by disruption of the *LDLR* negative regulator *MYLIP* (Fig 2B, Supplemental Tables 1-2). Using a stringent cutoff for screen analysis (false discovery rate, or FDR < 5%), we identified no other genes with a significant role in Lp(a) uptake. Using a more liberal significance threshold (FDR < 20%), we identified another 8 genes whose disruption was associated with decreased Lp(a) uptake. This group included the canonical *LDLR* regulators *SCAP* and *MBTPS2* and was significantly enriched for gene-gene interactions and functional annotations related to cholesterol metabolism (Fig S3). Among genes not classically associated with LDLR regulation but identified in our prior analogous screen of LDL uptake, we observed a significant correlation between each gene’s influence on LDL and Lp(a) uptake (Fig 2C, *r* = 0.63, p < 0.001). In a focused analysis of specific receptors previously proposed to mediate Lp(a) uptake, we did not identify a significant effect in our screen for any aside from *LDLR* (Fig 2D).

### LDLR expression modulates HuH7 Lp(a) uptake

To validate the functional influence of *LDLR* on Lp(a) uptake by HuH7 cells, we next generated and analyzed HuH7 cells with intact or genetically disrupted endogenous *LDLR* in the presence or absence of heterologous *LDLR* cDNA expressed from a lentiviral construct (Fig 3A). Because of the potential for contaminating LDL in Lp(a) preparations to confer a false positive dependence on *LDLR*, in our validation experiments we measured Lp(a) uptake with a secondary method of detection that was specific for Lp(a) (an ELISA for the apo(a) component of Lp(a) that is absent in LDL particles). To exclude the possibility of the fluorescent label mediating the interaction between Lp(a) and LDLR, we performed validation testing with native, unlabeled Lp(a) preparations. Consistent with our screen findings, we found that disruption of endogenous *LDLR* abrogated cellular Lp(a) uptake; this phenotype was not due to a CRISPR off-target effect since it was rescued by heterologous expression of *LDLR* cDNA (Fig 3B). Further supporting the role of *LDLR* in Lp(a) uptake, we also found that overexpression of *LDLR* on a wild-type background significantly augmented cellular uptake of Lp(a) (Fig 3B).

**Figure 3.**
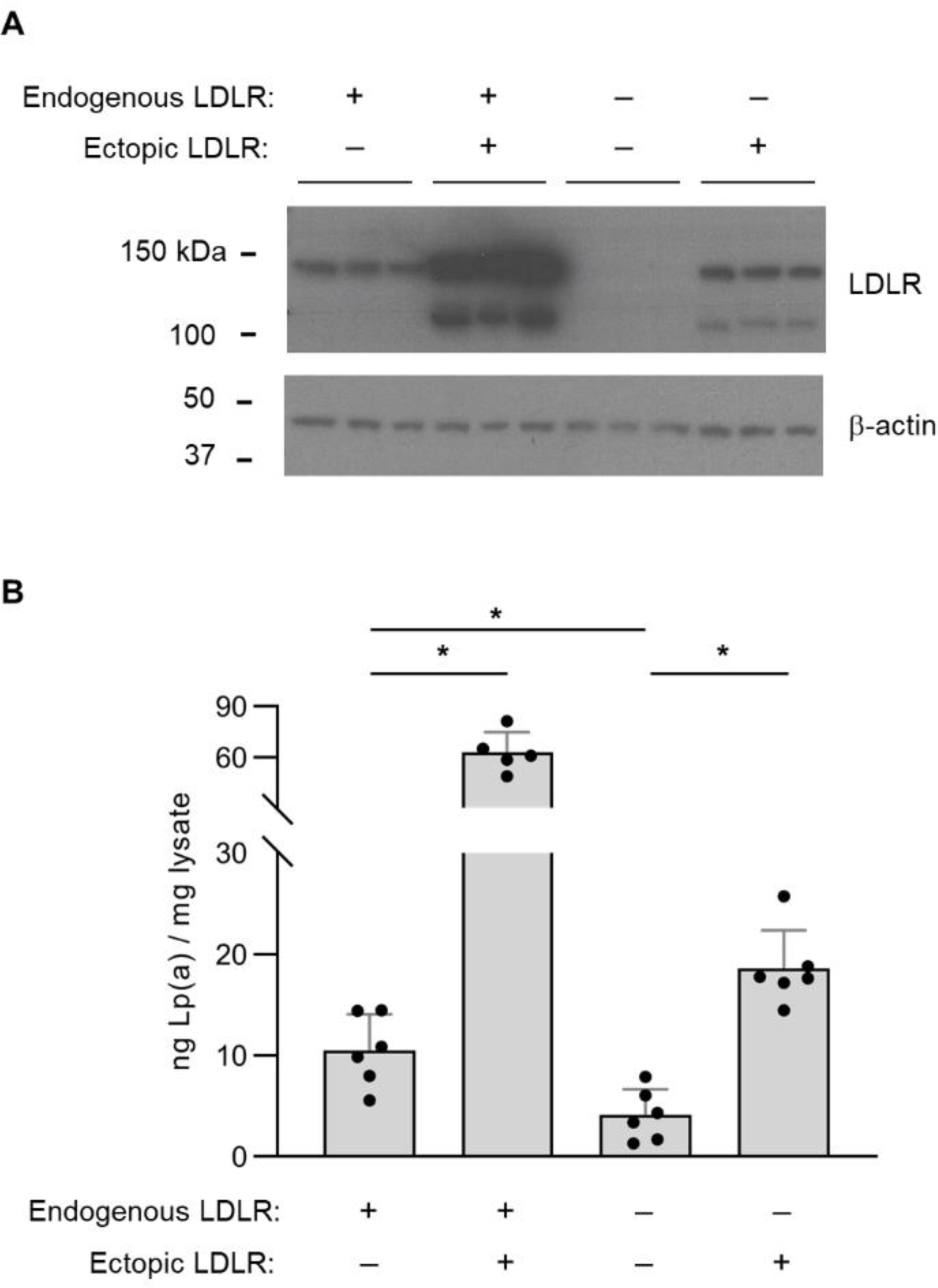
Analysis of Lp(a) uptake in *LDLR*-disrupted and *LDLR*-overexpressing HuH7 cells. (A) Immunoblot analysis of LDLR protein abundance in wild-type HuH7 cells and a clonal HuH7 cell line harboring a homozygous frameshift-causing insertion in exon 6 of *LDLR*, each with and without transduction of a lentiviral expression construct of a *LDLR* cDNA. (B) Quantification of cellular apolipoprotein(a) internalization for the cell lines indicated in (A) after incubation with 50 µg/mL Lp(a) for 3 hrs. Individual data points represent the mean of technical duplicates for each independent biologic replicate. Asterisks indicates p<0.05 by Student’s t-test and error bars depict standard deviation.

### Lp(a) directly binds to the LDL receptor extracellular domain

The functional influence of LDLR on HuH7 Lp(a) uptake might have been mediated by direct binding to Lp(a) or by an indirect effect of LDLR perturbation such as an alteration of cellular metabolism or the expression of other genes. To evaluate for the potential of Lp(a) to directly bind the LDL receptor, we used bio-layer interferometry, a label-free method in which binding to a ligand is detected by a change in the refractive index for white light^58^. We immobilized biotinylated recombinant LDLR extracellular domain to streptavidin-coated sensors and incubated with a range of concentrations of Lp(a) or the positive and negative controls LDL and HDL, respectively. As expected, we observed a concentration-dependent shift in refracted light for LDL with LDLR-immobilized sensors but not with control streptavidin-coated sensors alone, while parallel reactions with HDL revealed no evidence of LDLR binding (Fig 4A, C). As with LDL, Lp(a) exhibited a concentration-dependent and LDLR-dependent shift in refracted light, albeit of lesser magnitude than that observed for LDLR (Fig 4B, C).

**Figure 4.**
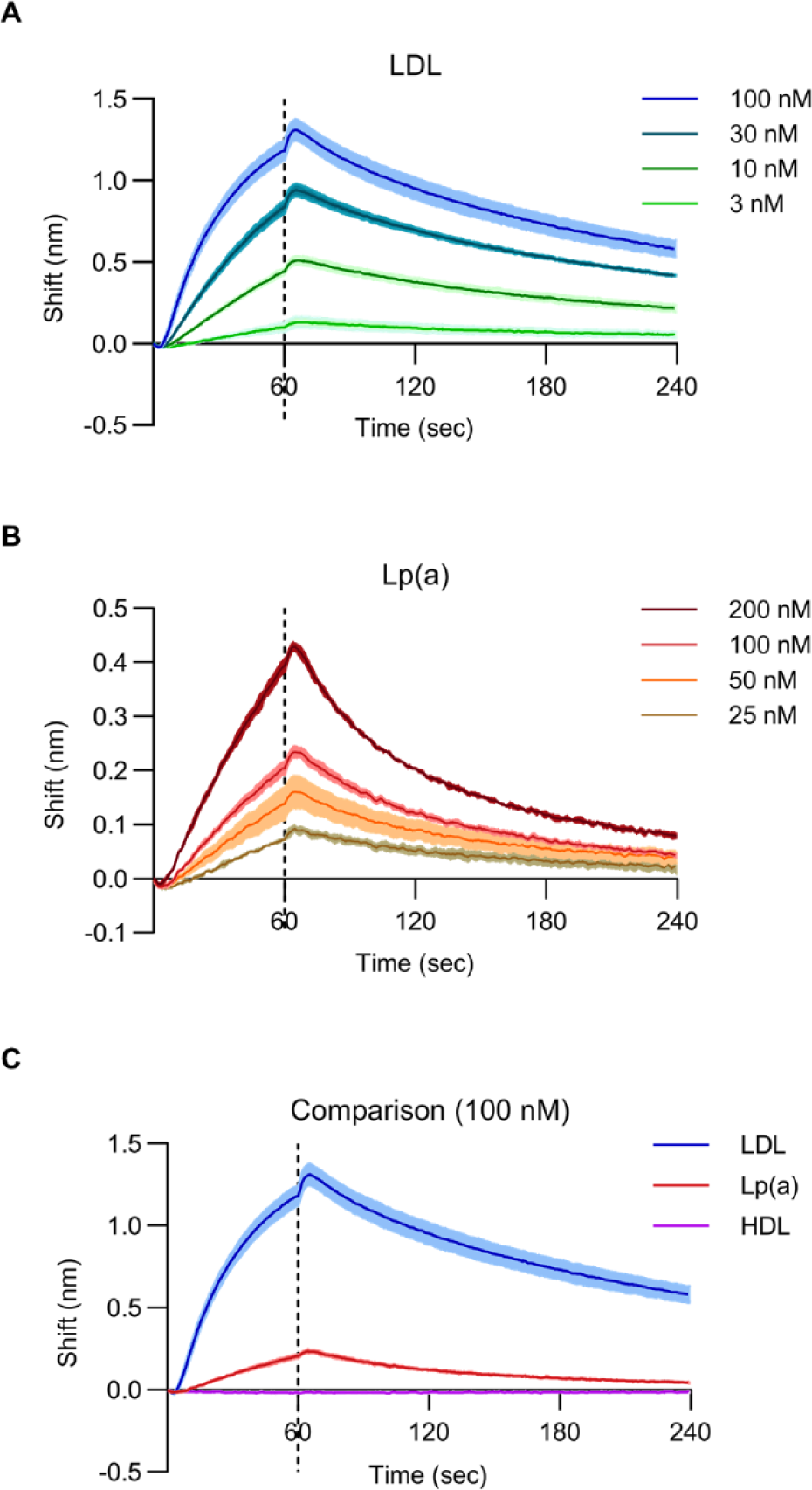
*In vitro* binding between Lp(a) and the LDLR extracellular domain. (A) Biolayer interferometry measurements for LDLR binding with LDL at the indicated concentrations. (B) Biolayer interferometry measurements for LDLR binding with Lp(a) at the indicated concentrations. (C) Comparison of biolayer interferometry measurements for LDLR binding to LDL, Lp(a), and HDL each at 100 nM. For all panels, solid lines indicate mean values and shaded regions indicate 1 standard deviation above and below the mean. Sensors were incubated with lipoprotein suspensions at t = 0 seconds and transferred to buffer only at t = 60 seconds, indicated by the dashed line.

### Humans with pathogenic *LDLR* alleles have elevated levels of circulating Lp(a)

To examine the physiologic relevance of our *in vitro* findings, we also analyzed Lp(a) levels and their relationship to *LDLR* genotype among individuals in the UK Biobank cohort^59^. We identified a total of 225 carriers of 25 different *LDLR* alleles with the highest level of evidence for pathogenicity as determined by an expert panel^60^. Each of these 225 individuals were heterozygous for the pathogenic allele. We adjusted lipoprotein levels for the use of lipid-lowering medications and controlled for age, sex, and ancestry (Supplemental Table 3). As expected, carriers of *LDLR* pathogenic alleles were found to have LDL cholesterol levels that were significantly greater than noncarriers (32.3% increase, p = 4.1 x 10^-116^, Fig 5A and Supplemental Table 4). Likewise, pathogenic *LDLR* allele carriers also exhibited significantly higher Lp(a) levels than noncarriers, though with a smaller magnitude of increase (14.3% increase, p = 1.6 x 10^-4^, Fig 5B and Supplemental Table 4). This relative increase in Lp(a) levels was statistically significant and of a comparable effect size whether analyzing raw or adjusted Lp(a) levels, untransformed or rank-based inverse-normalized values, all ancestries in the cohort or Europeans only, or with or without adjustment for genotype at the rs10455872 SNP that tags a narrower range of *LPA* KIV2 copy number variants and represents one of the top associations with Lp(a) levels in multiple genome-wide association studies^8,44–46^ (Fig S4 and Supplemental Table 4).

**Figure 5.**
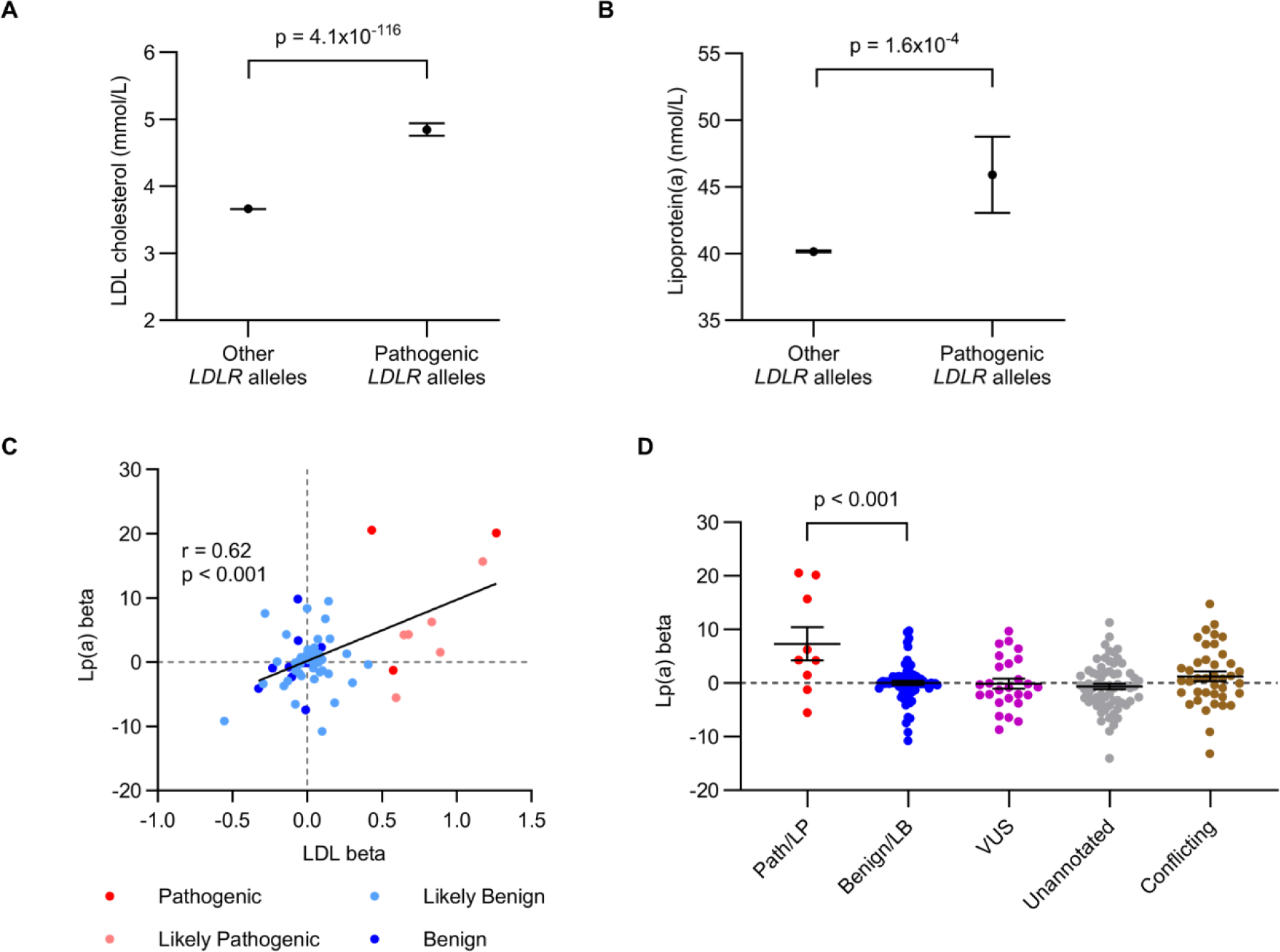
Analysis of Lp(a) levels for carriers of *LDLR* pathogenic alleles in the UK Biobank cohort. (A-B) Adjusted LDL and Lp(a) levels aggregated for 225 carriers and 358,405 noncarriers of 25 expert-curated pathogenic *LDLR* alleles in UK Biobank. Statistical analysis for (A) and (B) was performed with the burden testing framework in Regenie. (C) Correlation between LDL and Lp(a) beta coefficients for all single *LDLR* variants with the indicated annotations and a minor allele count of at least 25 in UK Biobank. Individual data points represent single variants and are colored by functional annotation. Regression analysis was performed with GraphPad Prism. (D) Lp(a) beta coefficients for single variants represented by at least 25 individuals in UK Biobank, grouped by ClinVar annotation with statistical analysis between groups performed by Student’s t-test assuming equal variance. For all panels, horizontal lines represent mean values and error bars indicate standard error of the mean.

We also analyzed the relationship of Lp(a) levels to a broader set of *LDLR* variants with different functional annotations (Supplemental Table 5). To avoid increased variability in small sample sizes, we focused our analysis on *LDLR* alleles represented by at least 25 carriers. We found that relative to alleles annotated as “Benign” or “Likely Benign”, those annotated as “Pathogenic” and “Likely Pathogenic” were associated with significantly increased Lp(a) levels (Fig 5C, p = 0<0.001). Among these single variants, there was a significant correlation (r = 0.64, p <0.0001) between the change in associated LDL and Lp(a) levels (Fig 5D). There was no significant increase in Lp(a) levels for carriers of *LDLR* variants lacking a functional annotation or annotated with “uncertain significance” or “conflicting interpretations of pathogenicity” (Fig 5C).

## DISCUSSION

A role for the LDL receptor in Lp(a) clearance has long been suspected due to the structural similarities between Lp(a) and LDL particles. Lp(a) is comprised of an LDL-like moiety which includes the LDLR ligand apolipoprotein B covalently bound to a single apo(a) molecule via disulfide linkage. Indeed, multiple studies in cells have provided support for an interaction between Lp(a) and LDLR^25–28^. However, other studies have found no influence of LDLR on cellular Lp(a) binding or uptake *in vitro*^29–32^. Likewise, while one study in humans found a defect in Lp(a) catabolism for patients with Familial Hypercholesterolemia (characterized by a defect in LDLR-mediated LDL uptake)^26^, other studies did not^35–37^. Although mice do not produce endogenous Lp(a), studies of their catabolism of exogenous human Lp(a) have also yielded mixed results, with some demonstrating LDLR-dependent^33,34^ and some LDLR-independent^23,30^ clearance. The basis for these discrepancies is unclear but may be related to differences in experimental design, including methods of Lp(a) purification and measurement. In this context, we now report the first forward genetic screen in human cells for cellular regulators of Lp(a) uptake, controlling for technical differences by functionally testing all potential Lp(a) receptors simultaneously. Overall, our screen results indicate a primary role for the LDL receptor in mediating cellular Lp(a) uptake under these conditions, with *LDLR* and its negative regulator *MYLIP* representing the top hits influencing Lp(a) uptake and no other potential receptor demonstrating a significant effect.

In addition to the functional data supporting a role for LDLR in Lp(a) uptake, we also found evidence of direct binding between Lp(a) and the LDLR extracellular domain. The magnitude and affinity of Lp(a) binding was qualitatively less than we observed for LDL binding, consistent with prior studies that reported reduced binding of Lp(a) relative to LDL for primary human hepatocytes, HepG2, and Hep3B cells^61,62^. Although our findings indicate direct binding between Lp(a) and LDLR *in vitro*, we cannot conclude whether this interaction is sufficient for physiologically relevant binding *in vivo* or if cooperative interactions with other molecules at the cell surface might have a role in increasing the affinity of Lp(a)-LDLR binding.

Prior human genetic studies on the association between *LDLR* alleles and Lp(a) levels have also produced conflicting results. Some studies of individuals with Familial Hypercholesterolemia (FH) have reported an increase in Lp(a) levels^38–40^, but others have not^41–43^ and suggested that the former may have been confounded by diagnostic ascertainment bias and/or unequal distributions of *LPA* risk alleles between FH and non-FH populations^42,43^. Multiple genome-wide association studies (GWAS) did not find an association between Lp(a) levels and common variants at the *LDLR* locus^8,44,45^, but a recent GWAS of 371,212 individuals in the UK Biobank cohort did^46^. Analyzing the same cohort, we now report an association between rare pathogenic *LDLR* alleles and increased Lp(a) levels, both by an aggregate analysis of 25 expert-curated pathogenic alleles with the highest level of evidence and by single variant analysis of a broader set of *LDLR* alleles. However, we found a smaller effect on Lp(a) than LDL levels, with the former increased by ∼14% and the latter ∼32% in pathogenic *LDLR* allele carriers. This finding is consistent with our lipoprotein uptake studies that revealed a less pronounced defect in *LDLR*-disrupted cells for Lp(a) uptake (∼61% reduction) compared to LDL uptake (∼87% reduction^54^), and with our *in vitro* binding studies showing a reduced affinity for LDLR binding to Lp(a) relative to LDL. It is also possible that the aggregate effect of these pathogenic *LDLR* alleles on Lp(a) may be blunted by the inclusion of individual variants with discordant effects on LDL and Lp(a) uptake. Further investigations will be necessary to establish the structural basis of the Lp(a)-LDLR interaction and its sensitivity to *LDLR* variants and different Lp(a) components including apo(a).

Modulation of LDLR activity has been a highly successful strategy for LDL-lowering and the prevention and treatment of atherosclerotic cardiovascular disease. Paradoxically, while PCSK9 inhibitors (which upregulate LDLR) have been found to significantly lower Lp(a), statins (which also upregulate LDLR by a different mechanism) do not^63^. A variety of models have been proposed to reconcile these findings, including the possibilities that Lp(a)-lowering by PCSK9 inhibitors may be mediated by receptors other than LDLR^64^, by an effect on Lp(a) synthesis rather than clearance^32,65^, or by a dependence of Lp(a) clearance on the expression level of LDLR and the relative concentrations of LDL and Lp(a)^66,67^. Intriguingly, a recent study also found that statin treatment caused a significant increase in *LPA* expression in HepG2 cells^17^. If the same is true for hepatocytes *in vivo*, it is possible that this concurrent Lp(a)-increasing effect of statins may offset and mask any Lp(a)-lowering resulting from increased LDLR-mediated clearance.

There are intrinsic limitations to our CRISPR screening approach that warrant consideration. First, a gene knockout screen such as ours may not detect an effect for essential genes, as cells harboring gRNAs targeting these genes are expected to become progressively depleted in pooled cultures prior to selection. Second, although HuH7 cells have been shown to serve as excellent models for lipoprotein uptake by hepatocytes^68^, the expression of any single gene may vary from its *in vivo* state in hepatocytes, potentially leading to a false negative result if a putative Lp(a) receptor is poorly expressed. Third, although the GECKOv2 library is optimized for activity and tests multiple gRNAs per target gene, stochastic variation in gRNA efficiency may lead to inefficient editing for a given target that could similarly cause a false negative result. Finally, loss-of-function CRISPR screens have limited power to detect a role for functionally redundant genes. For example, if a Lp(a) receptor were encoded by two different paralogues, then the intact function of one gene may compensate for loss of the other. These caveats notwithstanding, our screen findings together with our subsequent validation testing, *in vitro* binding studies, and human genetic analysis all lend support to the central importance of LDLR in mediating Lp(a) uptake by hepatocytes.

## Acknowledgments

This research was supported by the National Institutes of Health K08-HL148552 (BTE), R01-HL167733 (BTE), R35-GM127303 (AVS), R01-DK128871 (EKS), and R01DK131787 (EKS), the University of Michigan MBioFAR award (EKS), and the A. Alfred Taubman Medical Research Institute (BTE).

## Author Contributions

TGK and BTE conceived the project. TGK, JBC, MB, and CR performed the experiments. All authors contributed to data analysis and interpretation. TGK, JBC, and BTE wrote the manuscript with input from all authors. All authors reviewed final version of the manuscript.

## MATERIALS AND METHODS

### Cell lines and reagents

HuH7 and HEK-293T cells (ATCC, Manassas VA) were cultured in DMEM supplemented with 10% fetal bovine serum, 10 U/mL penicillin, and 10 µg/mL streptomycin (ThermoFisher Scientific, Waltham MA) in a humidified 5% CO2 chamber at 37°C. Cell lines were periodically tested for mycoplasma contamination and verified by microsatellite genotyping. Generation and genotyping of a *LDLR*-disrupted HuH7 clonal line was previously described^54^. A *LDLR* expression construct was prepared by PCR amplification of *LDLR* cDNA from a plasmid template (GenScript Biotech, Piscataway NJ, #OHu22799) with primers (Integrated DNA Technologies, Coralville IA) designed to provide flanking homology arms for HiFi assembly (New England Biolabs, Ipswich MA) into EcoRI/NotI-digested (New England Biolabs) LeGO-ACE2-IRES-blast^69^. Lentivirus was generated as previously described^55^ and used to either overexpress or rescue *LDLR* by transduction of HuH7 wild-type or *LDLR*-disrupted cells, respectively. Lipoprotein preparations were purified from human plasma, validated against lipoprotein standards by SPIFE Vis Cholesterol assay (Helena Laboratories, Beaumont TX, #3218), and provided by a commercial supplier (Athens Research and Technology, Athens GA, #12-16-121601, #12-16-120412, #12-16-080412).

### Lp(a) labeling

For initial pilot studies, purified Lp(a) preparations were treated with a variety of fluorescent labeling strategies – FITC Conjugation Kit (Abcam, Cambridge UK, #ab188285), Dil (1,1’-Dioctadecyl-3,3,3’,3’-Tetramethylindocarbocyanine Perchlorate) Stain (Thermo Fisher Scientific, #D282), and pHrodo Red Microscale Labeling Kit (Thermo Fisher Scientific, #P35363) – all per manufacturer’s instructions. Subsequent labeling experiments, including for the genome-scale CRISPR screen, were performed using pHrodo iFL Red Antibody Labeling Kit (Thermo Fisher Scientific, #P36014) at a lipoprotein to dye molar ratio of 1:28 for 15 mins in the dark at room temperature. Unconjugated free dye was removed from each sample by extensive dialysis against PBS. A portion of labelled and unlabeled lipoprotein samples were analyzed by SDS-PAGE on NuPAGE 3-8% gradient Tris-Acetate gels (Thermo Fisher Scientific, # EA0375) under reducing and nonreducing conditions followed by SilverQuest Silver Staining (Thermo Fisher Scientific, # LC6070) per manufacturer’s instructions or by in-gel fluorescence scanning with settings for Red Epifluorescence excitation and 700/50 nm emission using a ChemiDoc MP Imaging System (Bio-Rad Laboratories, Hercules CA).

### Analysis of Lp(a) uptake

Cells were seeded in 6-well or 12-well plates and analyzed 2-3 days later when their confluence reached 70-90%. To quantify cellular Lp(a) uptake, monolayers were washed once with serum-free DMEM and incubated in serum-free DMEM containing unlabeled or fluorescent Lp(a) at the indicated concentration for the indicated duration at 37°C. For flow cytometry analysis, cells were detached with TrypLE express (Thermo Fisher Scientific), washed twice with ice cold PBS, resuspended in ice-cold PBS, and analyzed on a Ze5 flow cytometer (Bio-Rad Laboratories) with gating and quantification of fluorescence intensity performed using FlowJo. For apo(a) ELISA, cells were washed once with PBS, scraped into ice-cold PBS suspensions, and centrifuged at 500 x g. Protein lysates were extracted from cell pellets by resuspension and incubation in RIPA buffer with end-over-end rotation for 15 mins followed by centrifugation at 21,000 x g for 30 min at 4°C and transfer of supernatants to new tubes for immediate analysis or storage at -80°C. Total protein concentrations of lysates were quantified by BCA Protein Assay (Thermo Fisher Scientific, #23225) and each sample was analyzed in technical duplicates for apo(a) protein concentration by ELISA (Abcam, #ab108878) with detection on a Spark Multimode Microplate Reader (Tecan, Männedorf Switzerland). Apo(a) concentrations were normalized to total protein concentrations of lysate and adjusted with subtraction of nonspecific background signal for control samples analyzed in parallel in the absence of exogenous Lp(a).

### CRISPR screen

For each independent biological replicate, a total of ∼60 million HuH7 cells distributed in 10 separate 15 cm^2^ cell culture plates at 70-90% confluence were transduced with a pooled lentiviral library containing the GeCKOv2 library^57^ at MOI ∼0.5. Selection of transduced cells with 2.5 μg/mL puromycin was started 24 hr post-transduction and maintained until no viable cells remained in parallel control plates that were not transduced with lentivirus. Cells were passaged every 2-3 days to maintain logarithmic phase growth with a minimum of ∼30 million cells, representing >200X library coverage, at all times. On day 14 post-transduction, pools of edited cells were incubated with 20 µg/mL pHrodo iFL Red-labelled Lp(a) in serum-free DMEM for 2 hrs at 37°C and prepared for flow cytometry as described above. Cell sorting was performed on a FACSAria III instrument (Becton Dickinson, Franklin Lakes NJ) with collection of 10% subpopulations of edited cells with the least and greatest magnitude of Lp(a) fluorescence.

Genomic DNA was extracted from sorted populations using a DNeasy kit (Qiagen, Hilden, Germany). A range of 5 to 9 million cells were collected in each bin for each replicate. Amplicon sequencing libraries were prepared as previously described^54,70,71^ and sequenced with a 1 x 75 single end read on a NextSeq instrument (Illumina, San Diego CA). Individual gRNA reads were extracted from FASTQ files with PoolQ and analyzed for enrichment using MAGeCK^72^. Genes identified as Lp(a) uptake modifiers with FDR<20% were analyzed for enrichment in Gene Ontology annotations using PANTHER v18.0^73^ and for gene-gene interactions using STRING v12.0^74^ with visualization of results by Cytoscape v3.9.1^75^.

### *In vitro* binding assays

Biolayer interferometry experiments were performed by incubating Octet Streptavidin Biosensors (Sartorius, Göttingen Germany, #18-5019) with 10 ug/mL biotinylated recombinant LDLR extracellular domain (BPS Bioscience, San Diego CA, #71206) and transferring LDLR-immobilized or control sensors to multiwell plates containing the indicated lipoprotein concentration in Octet Kinetics Buffer (Sartorius, 1#8-1105) at room temperature for 60 seconds followed by transfer to wells containing buffer only for an additional 180 seconds. Detection was performed with measurement of light interference every 0.2 seconds for 4 minutes on an Octet RED96 System instrument (Forte Biosciences, Dallas TX).

### UK Biobank analysis

The UK Biobank has been previously described^59^. Protocols were approved by the North West Multi-centre Research Ethics Committee and analyses in this project were conducted under UKBB Resource Project 18120 with IRB approval. Variant calling and mapping of the July 2022 release of whole exome sequencing (WES) was performed by the OQFE procedure as described by Krasheninina et al^76^. Quality control of variant call-rate, sample call-rate, read-depth, allelic balance, and Hardy-Weinberg equilibrium was performed as described by Szustakowski et al^77^. *LDLR* variants were selected with Annovar^78^ using the hg38 refGene database. Analysis was performed controlling for age (Datafield 21003), age^2^, sex (Datafield 22001), and the first 10 genetic principal components (Datafield 22009). A parallel analysis was also conducted that included genotype at the rs10455872 SNP as a covariate. Genetic relatedness was accounted for using array genotypes^79^. Individuals were excluded if they did not contain WES, genotyping array, serum LDL, and serum Lp(a) data. Summary statistics regarding the study population can be found in Supplementary Table 3. A total of 359,090 individuals were considered for the final analysis. Direct LDL cholesterol (Datafield 30780) and Lp(a) levels (Datafield 30790) were initially extracted from both instance 0 and instance 1. The final serum levels per individual were chosen by selecting the earliest instance with both non-missing serum values. Age and medication use (Datafield 20003) were retrieved to match the instance used for the phenotype. LDL levels were adjusted by an increase of 25% for the use of lipid-lowering medications as previously described^80^. Lp(a) levels were adjusted with decrease by 10% for the use of statins or an increase of 30% for the use of niacin based on estimated effects of these medications^81^. An adjustment for PCSK9 inhibitors was omitted as no individuals in this data release were identified as taking these medications. A list of the included medications and their codes is listed in Supplemental Table 3. Burden testing was performed using a pre-defined set of 61 variants annotated in ClinVar^82^ (accession SCV004022429.1) as pathogenic based on Clinical Genome Resource (ClinGen) Familial Hypercholesterolemia Variant Curation Expert Panel consensus guidelines^60^. SNPs were tested using Regenie^79^. Association testing was conducted for raw, adjusted, and rank-based inverse normalized transformations of the LDL and Lp(a) levels. We also performed a parallel analysis of Europeans only as determined by Oliveri et al^83^. ClinVar functional annotations of individual variants were extracted from gnomADv4.0^84^.

---

**Figure S1.**
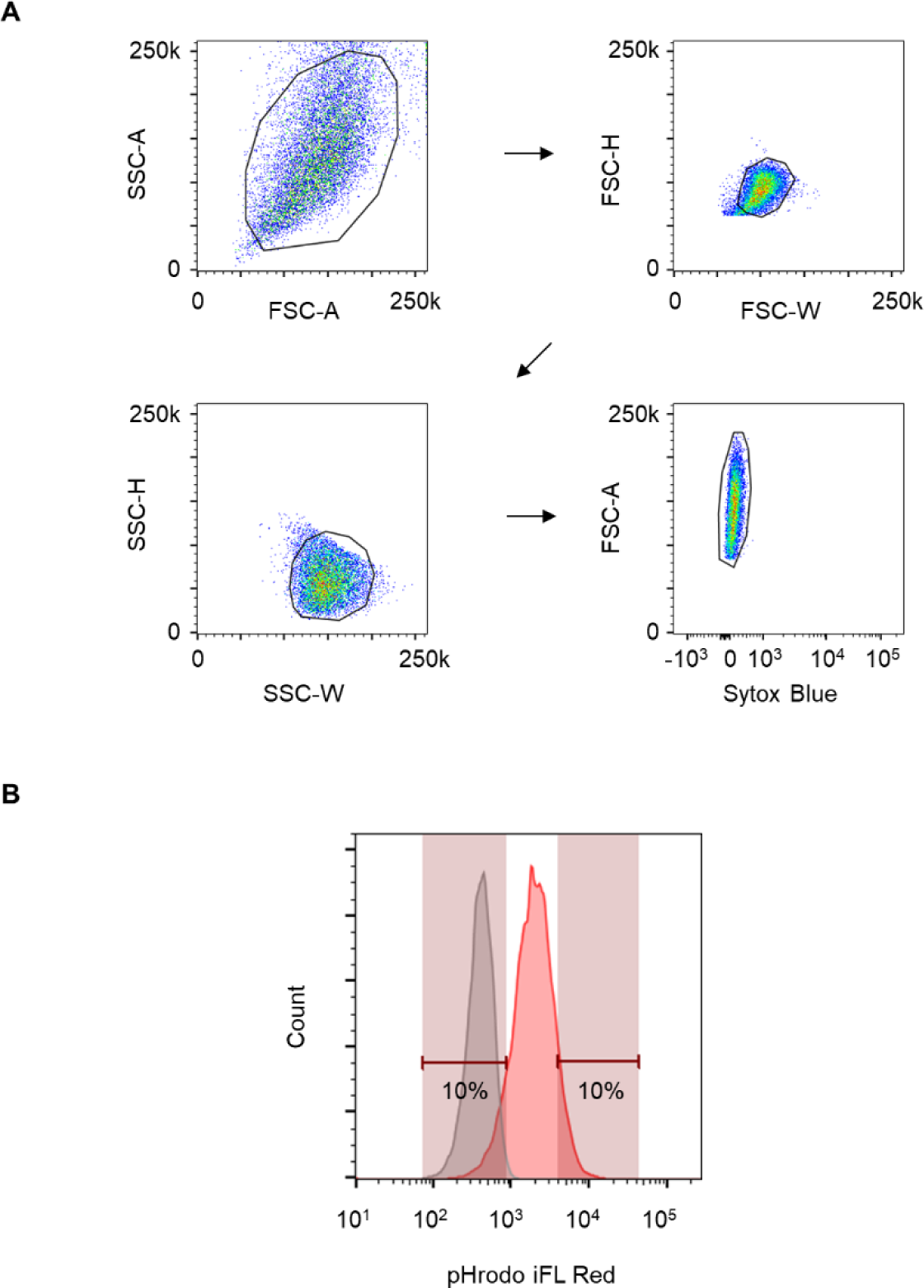
Detection of cellular uptake of labeled Lp(a) by flow cytometry. (A) Representative gating strategy for detection of HuH7 cells incubated with pHrodo iFL Red-labeled Lp(a). (B) Representative cell sorting strategy for the subpopulations of HuH7 cells with the greatest and least uptake of labeled Lp(a). Gray and red histograms represent cells incubated in the absence or presence of labeled Lp(a), respectively.

**Figure S2.**
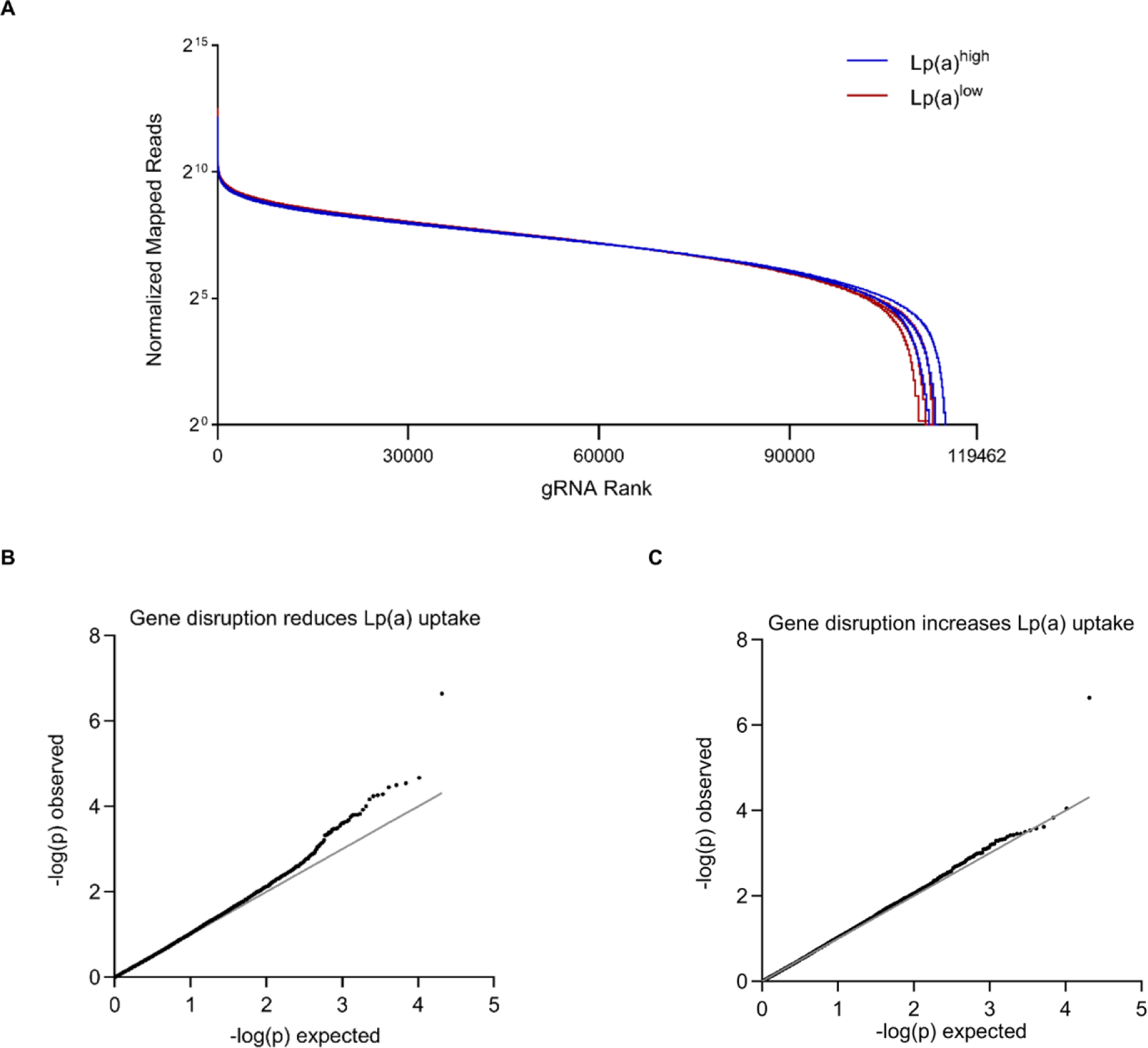
CRISPR screen quality control. (A) Cumulative distribution functions of normalized read counts for each gRNA in Lp(a)^high^ and Lp(a)^low^ populations for each of 3 independent biologic replicates. (B-C) Q-Q plots of observed versus expected –log(p-value) for gene perturbations associated with decreased (B) or increased (C) Lp(a) uptake. Observed p-values were calculated by MAGeCK (Supplemental Table 1).

**Figure S3.**
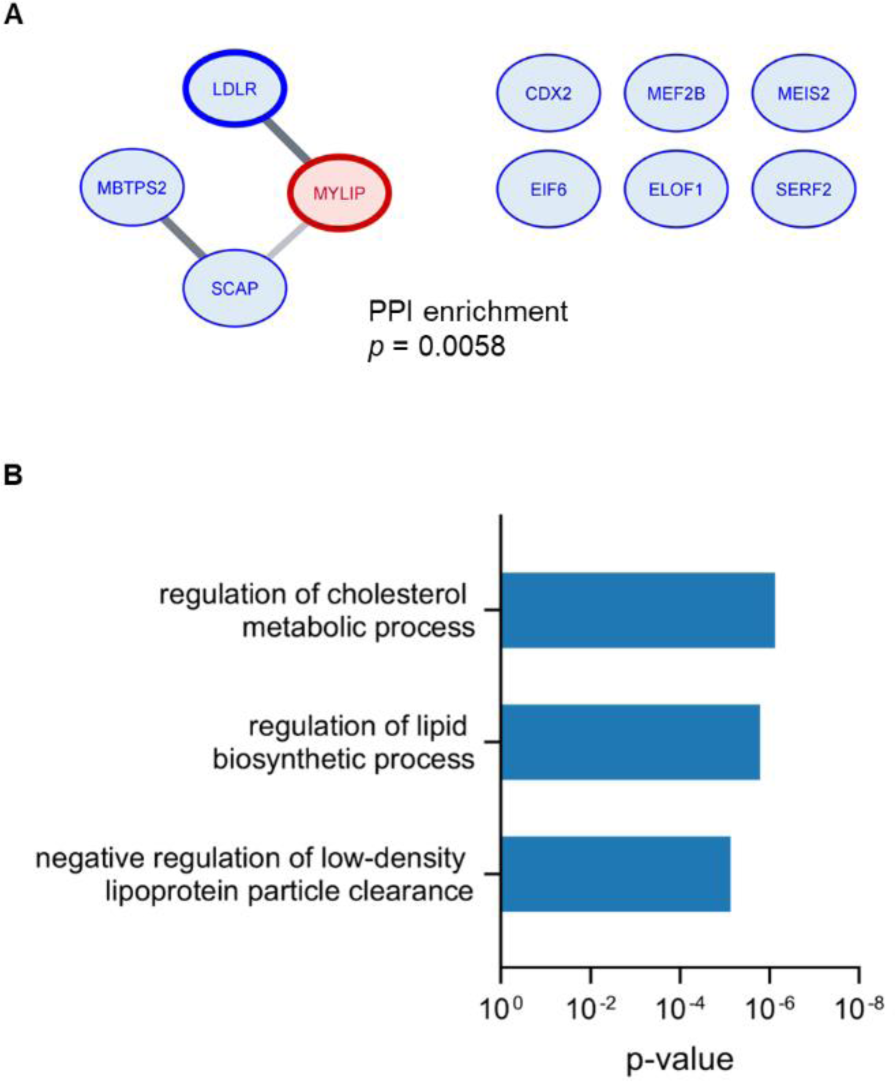
CRISPR screen analysis. (A) Network analysis of Lp(a) uptake modifiers identified in the genome-wide CRISPR screen with FDR < 20%. The borders of individual nodes are weighted by each gene’s −log(RRA score), and the lines connecting nodes are weighted by the strength of the protein–protein interaction within the STRING database. Genes whose disruption was associated with a decrease or increase in Lp(a) uptake are shaded in blue or red, respectively. The significance of the number of detected interactions relative to a randomly selected gene set was calculated by STRING. (B) Gene Ontology annotations for biologic processes enriched among the 10 genes identified in the Lp(a) uptake screen with FDR < 20% relative to the human genome. When multiple annotations within the same hierarchy were identified, the annotation with the most significant enrichment was selected for display. Calculation of p-values was performed by Fisher’s exact test using PANTHER.

**Figure S4.**
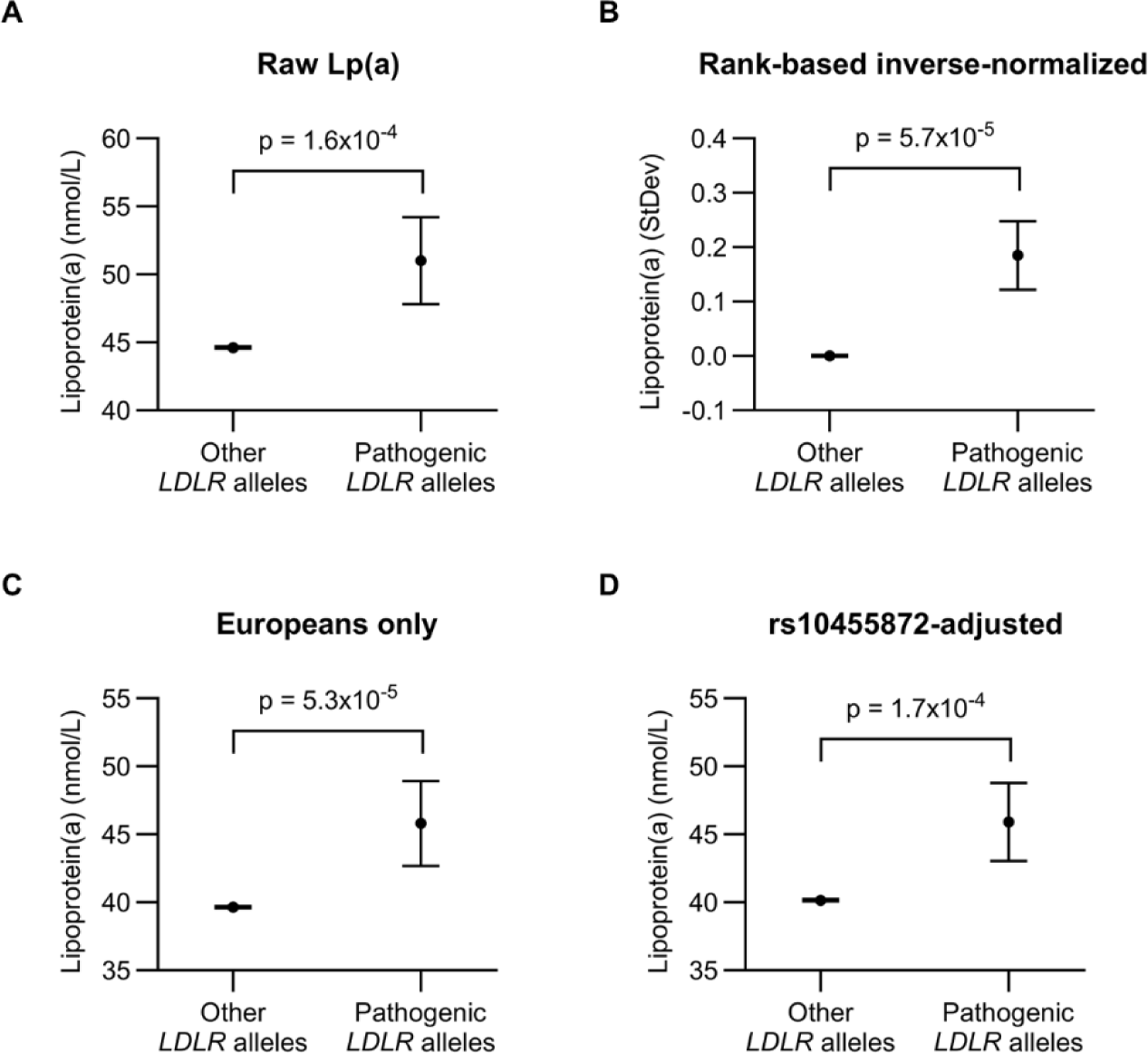
UK Biobank analysis of Lp(a) levels by *LDLR* genotype. Comparison of Lp(a) levels between carriers and noncarriers of 25 pathogenic *LDLR* alleles, as performed in Fig 5 with the following variations. (A) Comparison of raw Lp(a) levels. (B) Comparison of rank-based inverse-normalized adjusted Lp(a) levels. (C) Comparison of adjusted Lp(a) levels for the subset of individuals of European ancestry. (D) Comparison of adjusted Lp(a) levels with inclusion of rs10455872 genotype as a covariate.

**Supplemental Table 1.** Gene-level CRISPR screen results for modifiers of HuH7 cellular Lp(a) uptake. MAGeCK analysis of aggregate gene-level gRNA-level enrichment or depletion in Lp(a)^high^ relative to Lp(a)^low^ cells. Positive log2 fold change and RRA scores correspond to a gene’s disruption decreasing Lp(a) uptake; negative values correspond to a gene’s disruption increasing Lp(a) uptake.

**Supplemental Table 2.** Individual gRNA-level CRISPR screen results for modifiers of HuH7 cellular Lp(a) uptake. MAGeCK output for individual gRNA-level enrichment or depletion in Lp(a)^high^ relative to Lp(a)^low^ cells. Positive log2 fold change and RRA scores correspond to a gene’s disruption decreasing Lp(a) uptake; negative values correspond to a gene’s disruption increasing Lp(a) uptake.

**Supplemental Table 3.** Study characteristics of UK Biobank cohort. Demographic data, lipoprotein levels, and medication usage by individuals in the UK Biobank analyzed in this study.

**Supplemental Table 4.** Aggregate analysis of LDL and Lp(a) levels for carriers of 25 different pathogenic LDLR alleles. . LDL and Lp(a) analysis for carriers of 25 expert-curated pathogenic *LDLR* alleles, individually and in aggregate. Different tabs include data for raw lipoprotein levels, adjusted lipoprotein levels, rank based-inverse normalized values, and adjusted lipoprotein levels accounting for rs10455872 genotype as a covariate.

**Supplemental Table 5.** Single variant analysis of LDL and Lp(a) levels according to functional annotation. LDL and Lp(a) analysis for carriers of *LDLR* variants of the indicated functional annotations and allele frequencies in the UK Biobank cohort.

